## Supplementary material for "Full factorial construction of synthetic microbial communities": Source code 1 - protocol_generator.R

```
# customizable parameters
L <- 8 # landscape size (# of species; this is called m in the manuscript)
v0 <- 25 # minimum working volume (in uL)

# print warning
# print(paste('This will require ', ceiling(2^L/96), ' 96-well plates and 8 falcon tubes:', sep = ''))
# print(paste('   Each plate will hold ', L*v0, ' uL per well', sep = ''))
# print(paste('   Each tube will hold ', 3*10*v0*ceiling(2^L/96)/1000, ' mL', sep = ''))
# print('')

### FUNCTIONS

# binary representation of integer n (returns a string)
binary <- function(n) sapply(n, FUN = function(n) {
  binary_str <- ""
  while (n > 0) {
    if (n %% 2 == 1) {
      binary_str <- paste("1", binary_str, sep="")
    } else {
      binary_str <- paste("0", binary_str, sep="")
    }
    n <- bitwShiftR(n, 1)
  }
  if (binary_str == "") {
    binary_str <- "0"
  }
  return(binary_str)
})

# add leading zeros to binary number (as string)
add_leading_zeros <- function(n, length.out) as.character(sapply(n, FUN = function(n) {
  if (nchar(n) >= length.out) {
    return(n)
  } else {
    num_zeros <- length.out - nchar(n)
    return(paste(c(rep("0", num_zeros), substr(n, 1, length.out - num_zeros)), collapse = ''))
  }
}))

# returns n given its binary representation (as a string)
# this has a precision limit of 2^30 (will only work for landscapes of 30 species or less, but that protocol would require over 1e7 96-well plates anyway)
binary_to_integer <- function(binary_str) sapply(binary_str, FUN = function(binary_str) {
  n <- 0
  for (i in 1:nchar(binary_str)) {
    bit <- substr(binary_str, i, i)
    if (bit == "1") {
      n <- n + 2^(nchar(binary_str)-i)
    }
  }
  return(n)
})

# length of binary representation of integer n
binLength <- function(n) sapply(n, FUN = function(n) {
  length <- 0
  while (n > 0) {
    length <- length + 1
    n <- bitwShiftR(n, 1)
  }
  return(length)
})

# number of 1's in binary expansion of integer n ("binary weight")
binWeight <- function(n) sapply(n, FUN = function(n) {
  count <- 0
  while (n > 0) {
    if (n %% 2 == 1) {
      count <- count + 1
    }
    n <- bitwShiftR(n, 1)
  }
  return(count)
})

# does species sp go in column n? (n>1)
species_in_column <- function(n, sp) as.numeric((n-1) %% (2^(sp-3)) >= 2^(sp-4))

# how many parts are there in column n?
how_many_parts <- function(n) 3 + binWeight(n - 1)

# plates layout
n_plates <- ceiling(2^L/96)

where_species <- data.frame(species_in_column = character(0),
                            column_index = numeric(0))
for (sp in 4:L) {
  where_species <- rbind(where_species,
                         data.frame(species_in_column = sp,
                                    column_index = which(species_in_column(1:(2^L/8), sp) == 1)))
}

layout <- data.frame(column_index = 1:(2^L/8),
                     plate = paste('plate_', rep(1:n_plates, each = 12)[1:(2^L/8)], sep = ''),
                     column_in_plate = rep(1:12, ceiling(2^L/96))[1:(2^L/8)],
                     volume_water_fill = L - how_many_parts(1:(2^L/8)))
layout <- merge(layout, where_species, all = T)
layout$species_in_column[is.na(layout$species_in_column)] <- 'none'


### MAKE PROTOCOL

# step 1: fill plates with necessary water volume
protocol <- vector(mode = 'list')
protocol[['water_fill']] <- unique(layout[, c('volume_water_fill', 'plate', 'column_in_plate')])
protocol[['water_fill']] <- protocol[['water_fill']][order(as.numeric(gsub('plate_', '', protocol[['water_fill']]$plate)), protocol[['water_fill']]$column), ]
protocol[['water_fill']] <- protocol[['water_fill']][order(protocol[['water_fill']]$volume_water_fill, decreasing = T), ]
protocol[['water_fill']]$volume_water_fill <- v0 * protocol[['water_fill']]$volume_water_fill
colnames(protocol[['water_fill']]) <- c('volume_uL', 'plate', 'column')
protocol[['water_fill']] <- protocol[['water_fill']][protocol[['water_fill']]$volume_uL != 0, ]
rownames(protocol[['water_fill']]) <- NULL

# steps 2 to L-2: add species 4 to L to necessary columns
for (s in 4:L) {
  
  step_name <- paste('species_', s, sep = '')
  protocol[[step_name]] <- layout[layout$species_in_column == s, c('species_in_column', 'plate', 'column_in_plate')]
  protocol[[step_name]] <- cbind(volume_uL = v0, protocol[[step_name]][, 2:3])
  colnames(protocol[[step_name]]) <- c('volume_uL', 'plate', 'column')
  protocol[[step_name]] <- protocol[[step_name]][order(as.numeric(gsub('plate_', '', protocol[[step_name]]$plate)), protocol[[step_name]]$column), ]
  rownames(protocol[[step_name]]) <- NULL
   
}

# almost final step: assemble 3-species landscape
land3 <- add_leading_zeros(binary(0:7), 3)
layout_3 <- data.frame(species = character(0),
                       tube = numeric(0))

v0_tubes <- ceiling((2000 + 2^(L-3)*v0)/1000)*1000

protocol[['tubes_water_fill']] <- data.frame(volume_uL = v0_tubes*(3 - binWeight(0:7)),
                                             tube = paste('tube_', LETTERS[1:8], sep = ''))

for (s in 1:3) {
  
  step_name <- paste('tubes_species_', s, sep = '')
  protocol[[step_name]] <- data.frame(volume_uL = v0_tubes,
                                      tube = paste('tube_', LETTERS[as.numeric(which(sapply(land3, FUN = function(n) substr(n, 4-s, 4-s) == '1')))], sep = ''))
  
}


### WRITE UP PROTOCOL

how_much_monoc <- ceiling(v0*2^(L-1)/1000)*1000
how_much_water <- ceiling(L*v0*2^(L-1)/1000)*1000

# create output file (delete existing one if appropriate)
file_name <- './protocol.txt'
if (file.exists(file_name)) {
  file.remove(file_name)
}
file.create(file_name)

# write-up protocol
txt <- 'BEFORE WE START\n--------------------------------------------------------------------------------\n'
txt <- c(txt, paste('This protocol requires ', ceiling(2^L/96), ' 96-well plates and 8 falcon tubes:', sep = ''))
txt <- c(txt, paste('   Each plate will hold ', L*v0, ' uL per well', sep = ''))
txt <- c(txt, paste('   Each tube will hold ', 3*v0_tubes/1000, ' mL', sep = ''))
txt <- c(txt, paste('We need each of the ', L, ' species grown in monoculture (min. ', how_much_monoc/1000, ' mL of each monoculture, recommended ', 15 + how_much_monoc/1000, ' mL)', sep = ''))
txt <- c(txt, paste('We also need ', how_much_water/1000, ' mL of buffer (ddH2O, PBS, or carbon-free medium), recommended ', 15 + how_much_water/1000, ' mL.', sep = ''))
txt <- c(txt, 'In what follows, we will assume that the recommended starting volumes are available. If lower volumes are available, the third step of the protocol will need to be adjusted accordingly.\n')

txt <- c(txt, '\nSTEP 1: compensatory buffer to homogenize final densities\n--------------------------------------------------------------------------------\n')
txt <- c(txt, paste('Label ', n_plates,' 96-well plates as plate_1 to plate_', n_plates, sep = ''))
txt <- c(txt, 'Pipette the indicated volumes of buffer into the indicated columns:\n', '')
cat(paste(txt, collapse = '\n'), file = file_name)
suppressWarnings(write.table(protocol[['water_fill']], file = file_name, sep = '\t', row.names = FALSE, col.names = TRUE, quote = FALSE, append = TRUE))

txt <- '\n\nSTEP 2: inoculate monocultures\n--------------------------------------------------------------------------------\n'
txt <- c(txt, paste('Pipette the indicated volumes of species 4 to ', L, ' monocultures into the indicated columns:', sep = ''))
cat(paste(c(txt, ''), collapse = '\n'), file = file_name, append = TRUE)
for (i in 4:L) {
  cat(paste('\nSPECIES ', i, '\n', sep = ''), file = file_name, append = TRUE)
  suppressWarnings(write.table(protocol[[paste('species_', i, sep = '')]], file = file_name, sep = '\t', row.names = FALSE, col.names = TRUE, quote = FALSE, append = TRUE))
}

txt <- '\n\nSTEP 3: 3-species landscape\n--------------------------------------------------------------------------------\n'
txt <- c(txt, 'Label 8 Falcon tubes as tube_A to tube_H')
txt <- c(txt, 'Pipette the indicated volumes of buffer (ddH2O, PBS, or carbon-free medium) into the indicated tubes:\n')
cat(paste(c(txt, ''), collapse = '\n'), file = file_name, append = TRUE)
suppressWarnings(write.table(protocol[['tubes_water_fill']], file = file_name, sep = '\t', row.names = FALSE, col.names = TRUE, quote = FALSE, append = TRUE))
txt <- '\nPipette the indicated volumes of species 1 to 3 monocultures into the indicated tubes:'
cat(paste(c(txt, ''), collapse = '\n'), file = file_name, append = TRUE)
for (i in 1:3) {
  cat(paste('\nSPECIES ', i, '\n', sep = ''), file = file_name, append = TRUE)
  suppressWarnings(write.table(protocol[[paste('tubes_species_', i, sep = '')]], file = file_name, sep = '\t', row.names = FALSE, col.names = TRUE, quote = FALSE, append = TRUE))
}
cat('\nGently shake tubes to homogenize, then pipette:\n', file = file_name, append = TRUE)
for (i in 1:8) cat(paste('\n', 3*v0, ' uL per well from tube ', i, ' into row ', LETTERS[i], ' of plates 1 to ', n_plates, sep = ''), file = file_name, append = TRUE)

# final plate layout
txt <- '\n\n\nFINAL LAYOUT\n--------------------------------------------------------------------------------\n'
txt <- c(txt, 'Samples are located in the positions indicated below:')
cat(paste(c(txt, ''), collapse = '\n'), file = file_name, append = TRUE)
lout <- vector(mode = 'list',
               length = n_plates)
for (i in 1:n_plates) {
  plate_seq <- seq(96*(i-1), min(96*(i-1) + 95, 2^L - 1))
  plate_seq <- add_leading_zeros(binary(plate_seq), length.out = L)
  plate_seq <- c(plate_seq, rep('(empty)', 96 - length(plate_seq)))
  lout[[i]] <- matrix(plate_seq, nrow = 8, ncol = 12, byrow = FALSE)
  rownames(lout[[i]]) <- LETTERS[1:8]
  colnames(lout[[i]]) <- 1:12
  
  cat(paste('\nPLATE ', i, '\n', sep = ''), file = file_name, append = TRUE)
  suppressWarnings(write.table(as.data.frame(lout[[i]]), file = file_name, sep = '\t', row.names = TRUE, col.names = TRUE, quote = FALSE, append = TRUE))
  
}
```
